# Formation of the moving junction is the nexus for host cytoskeletal remodelling during *Plasmodium falciparum* invasion of human erythrocytes

**DOI:** 10.64898/2026.03.29.715162

**Authors:** Niall D Geoghegan, Cindy Evelyn, Aurelie Dawson, Danushka Marapana, Dawson B. Ling, Pradeep Rajasekhar, Michael J. Mlodzianoski, William Nguyen, Brad E. Sleebs, Christopher J. Tonkin, Lachlan W. Whitehead, Alan F. Cowman, Kelly L. Rogers

## Abstract

*Plasmodium falciparum* invasion of human erythrocytes is a complex and tightly coordinated process, involving host cell attachment, moving junction formation and engagement of the parasite’s actomyosin motor. The temporal precision of these events is mediated by distinct ligand-receptor interactions and the sequential release of the merozoite’s apical organelles. What remains unclear is how these molecular and biophysical interactions enable *Plasmodium* to bypass the stable erythrocyte membrane-cytoskeletal complex. Here, several *P. falciparum* lines expressing different fluorescently tagged apical organelle proteins, were imaged with lattice light sheet microscopy (LLSM) to determine the timing of cytoskeletal disassembly and apical organelle release. Blocking the AMA1-RON2 interaction has no effect on the PfRh5-basigin Ca^2+^ flux but prevents host cytoskeleton disassembly. In contrast, the inhibition of parasite actin polymerisation had no effect on cytoskeletal clearance but caused a sustained Ca^2+^ response. We further demonstrate that establishment of the moving junction is temporally linked to clearance of the host cytoskeleton.

Collectively, our findings support the existence of an association between the RON complex and components of the host cytoskeleton, which mediates the localised disruption of the erythrocyte-membrane cytoskeletal complex during invasion.

## INTRODUCTION

The asexual blood stage of the malaria causing parasite, *Plasmodium*, has a complex life cycle that includes invasion of the host erythrocyte by its merozoite form. The process of invasion is dynamic and involves a tightly coordinated series of molecular interactions ^1,2,3^, which can be broken into several key stages. The process begins with attachment of the merozoite to an erythrocyte via a variety of adhesins, released onto the parasite surface from apical organelles, called micronemes and exonemes, during parasite egress from an infected erythrocyte. These adhesins include the *P. falciparum* reticulocyte binding-like homologues (PfRh) and erythrocyte binding antigens (EBA) (reviewed in ^1^). This is followed by a period of deformation mediated by the merozoite during its initial interactions with the erythrocyte membrane. This culminates in apical reorientation through a process of host cell membrane wrapping ^4,5^ to position the apical end of the merozoite perpendicular to the plane of the erythrocyte membrane.

An essential protein complex released from the micronemes onto the merozoite surface includes PCRCR which is a pentameric complex made up of PTRAMP, CSS, Ripr, CyRPA and PfRh5. PTRAMP has a transmembrane region that anchors the complex to the merozoite membrane with PfRh5 at the tip enabling binding to the receptor basigin on the erythrocyte surface ^6–8^. PfRh5 – basigin binding is an essential ligand receptor interaction required for successful invasion that has been linked to a Ca^2+^ flux and the putative opening of a pore at the membrane through which the parasite can release its rhoptry contents ^3,9^. Within these contents is the RON-complex, which incorporates into the host erythrocyte membrane through its transmembrane protein PfRON2, which binds to PfAMA1 on the surface of the invading merozoite, forming a boundary between parasite and host cell, known as the moving junction (MJ) ^10^. One of the final steps is engagement of the merozoite’s actomyosin motor, that provides the force to propel the parasite through the MJ complex and into the newly forming parasitophorous vacuole (PV) ^11^.

The main physical barrier that the merozoite faces during erythrocyte invasion is the membrane-cytoskeleton complex of the erythrocyte. This complex is comprised of a lipid bilayer coupled to a pseudohexagonal mesh of spectrin tetramers by the actin junctional and ankyrin complexes ^12^. We have previously shown that the invading merozoite hijacks and reorganises the erythrocyte lipid bilayer to form the nascent parasitophorous vacuole, answering the long-standing question of whether the parasitophorous vacuole membrane (PVM) is host or parasite derived ^13^. What remains unknown, is how the parasite bypasses the host cytoskeleton barrier. During the period of active invasion, host cytoskeletal proteins (band3, actin, ankyrin and adducin) have been shown to be absent from the invasion site by fixed immunofluorescence imaging ^14^. In addition, at early stages of invasion, binding of the merozoite protein EBA175 to Glycophorin A (GPA) has been shown to induce phosphorylation of numerous cytoskeletal proteins and alter the biomechanics of erythrocytes ^15,16^. Similarly, using recombinantly expressed Rh5 it was found that ligation to its host receptor basigin induces the phosphorylation of proteins such as GPA, Glycophorin C (GPC) and ankyrin among others ^17^ further suggesting signalling events that occur in the erythrocyte upon parasite engagement. In addition to specific ligand receptor interactions, merozoite attachment alone can phosphorylate various cytoskeletal elements hinting at additional factors in this process ^14^. These findings combine to suggest a parasite driven disruption of the underlying host cytoskeleton; however, the precise timing of cytoskeletal breakdown remains unknown.

Following binding of the PCRCR complex to basigin it has been hypothesised that a pore forms, through which calcium (Ca^2+^) flows from the merozoite to the host erythrocyte ^3^. It has previously been hypothesised that this Ca^2+^ flux may play a role in the breakdown of the erythrocyte cytoskeleton ^18^. This process occurs during the recoil phase of the last deformation prior to merozoite internalisation and is localised to the parasite apex ^13,19^, however its function remains unknown.

Following this event, the rhoptry neck components are released and the RON complex, comprised of PfRON2, PfRON4 and PfRON5, is ejected into the host erythrocyte. PfAMA1, is bound to the merozoite membrane through its transmembrane region, and tightly binds to the RON complex via PfRON2 ^20^, also a transmembrane protein which inserts into the host erythrocyte membrane after it has been released from the rhoptry organelle. PfRON2 is part of a molecular assembly together with PfRON4 and PfRON5 ^21^, which are on the cytoplasmic side of the erythrocyte membrane ^20^. The RON complex is a key component of the MJ and is conserved across the apicomplexan phylum.

Many of the initial discoveries regarding the structure of the RON complex and its role in the establishment of the MJ were made in the apicomplexan parasite, *Toxoplasma gondii* ^22–25^. The RON complex forms a stable connection between invading parasite and host cell, potentially mediated through coupling to the underlying cytoskeleton ^25^.

Most studies interrogating changes to the erythrocyte cytoskeleton during *Plasmodium* invasion have been limited to the use of recombinant proteins or fixation of invading merozoites, both of which lack the temporal resolution to determine the highly dynamic sequence of steps leading to cytoskeletal breakdown. Lattice light-sheet microscopy (LLSM) has emerged as an effective tool to study the dynamics of host cell invasion owing to its high resolution in space and time and its capacity for 3-dimensional imaging ^6,13,26,27^. This technique was critical in understanding the dynamics of membrane reorganisation and PVM formation.

In this study, a comprehensive toolset was developed to image the key stages of invasion: deformation and reorientation, recoil, pore formation and Ca^2+^ flux, MJ formation and, finally, motor engagement and invasion. For the first time, using fluorescent parasite lines expressing combinations of PfRON3-mNeonGreen, PfRON4-mScarlet, and PfAMA1-mScarlet, formation of the MJ was directly imaged in the context of known invasion checkpoints. The breakdown of the host cytoskeleton correlates to the formation of the MJ, which then acts as a molecular sieve that sorts lipids and proteins into the forming PV. In contrast, the timing of the Ca^2+^ flux was not found to be tightly coupled to clearance of the host cytoskeleton, suggesting it is not involved in this process.

Furthermore, data presented here shows that the Ca^2+^ flux is prolonged in the presence of Cytochalasin D and *S*-W827, inhibitors of actin polymerisation, suggesting the parasite’s actomyosin motor plays a role in tuning the duration and decay of the Ca^2+^ flux observed during invasion.

## RESULTS

### A 4D spatio-temporal toolbox for understanding host cytoskeletal remodelling during *P. falciparum* invasion

Lattice light sheet microscopy (LLSM) is a gentle, live cell volumetric imaging method suitable for understanding the role of specific parasite ligand-host cell receptor interactions (e.g. PfRh5-Basigin), remodelling of the host membrane and formation of the nascent PV during invasion ^6,13,27^. To understand how *P. falciparum* interacts with the erythrocyte during invasion, an experimental toolbox was designed, comprising a range of fluorescent dyes and transgenic parasite lines expressing fluorescently tagged proteins to interrogate specific inhibitors of merozoite invasion. This toolbox was coupled with a LLSM platform, the Lattice Light Sheet 7 (LLS7, Carl Zeiss, Germany), which provides the means for gentle long-term imaging of invasion in a high throughput format and with more convenient glass bottom chamber slides (**Supplementary Fig. 1a (i-iv)**).

Using this approach, we were able to interrogate all stages of invasion, including the timing of rhoptry organelle discharge, formation of the MJ and when and how the host cytoskeleton breaks down (**Supplementary Fig. 1a (i-iv)**). Breakdown of the host cytoskeleton was visualised during invasion by labelling F-actin using a cell permeable fluorescent dye for live cell imaging. The fluorescence intensity and photostability of several commercially available dyes for labelling F-actin in human erythrocytes were firstly evaluated by LLSM (**Supplementary Fig 1b-e**). The far red fluorogenic, Silicon-Rhodamine, F-actin fluorescence dye, SiR-Actin (Spirochrome, 1 μM), was selected as providing the most optimal signal intensity over the background for the duration of an invasion experiment. Interestingly, a proportion of the SiR-Actin signal was maintained at steady state, despite a period of sustained photobleaching, which likely corresponded to the reservoir of dynamic actin filament formation in human erythrocytes ^28^. A range of cell permeable dyes were also used for membrane staining (e.g. Di-4-ANEPPDHQ, TF-cholesterol and TF-PC) and for parasite tracking (e.g. MitoTracker, Fluo-4 AM) (**Supplementary Fig. 1a (i)**).

Labelling of the erythrocyte boundary by SiR-Actin, provided the ability to visualise parasite induced deformations of the cell (**Fig. 1a, Supplementary Movie 1**). It was observed that the actin cytoskeleton remained intact when deformations of the host erythrocyte were made by merozoites attempting to invade, and that break down of the cytoskeleton occurred prior to and during internalisation by an invading parasite. When uninfected erythrocytes are also counterstained with the membrane dye, Di-4-ANEPPDHQ, the actin and membrane channels overlap for all parasite induced host cell membrane deformations preceding the step of internalisation. However, cytoskeletal clearing became distinguishable only during internalisation and PVM formation (**Fig. 1b, Supplementary Movie 1**).

**Figure 1:**
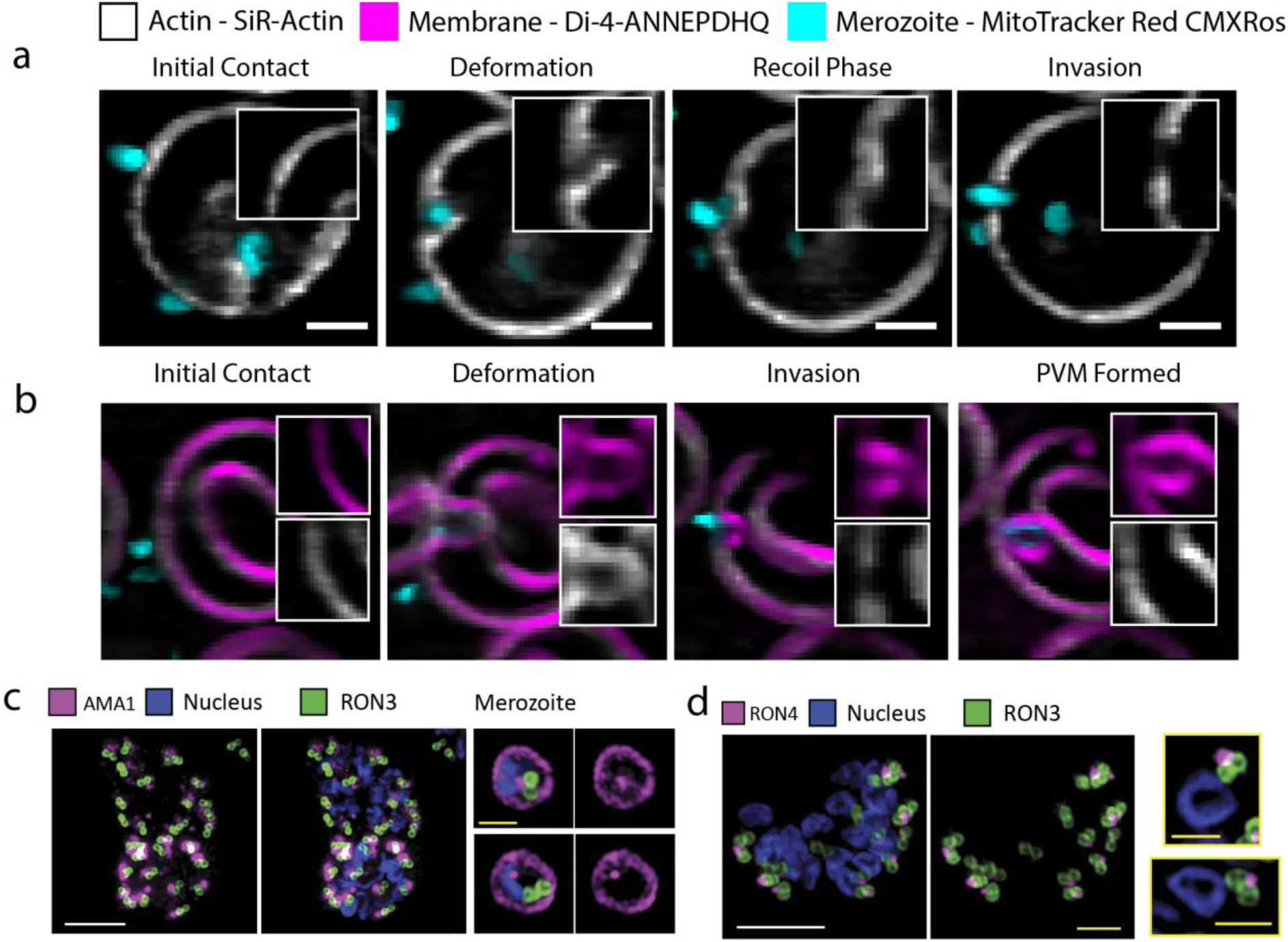
A spatio-temporal toolset to quantify invasion and formation of the moving junction. (a) Representative Lattice Light-sheet microscopy images showing different stages of invasion in the context of host cytoskeletal breakdown. Inset shows zoomed region of cytoskeletal breakdown through SiR-Actin labelling. Scale bar – 2 µm. (b) Representative LLSM images showing key invasion steps in the context of PVM formation and cytoskeletal breakdown together. Scale bar – 2 µm. 3D-SIM images of schizont and merozoite stage parasites were captured for (c) dual PfRON4-mScarlet/PfRON3-mNeonGreen, and (d) PfAMA1-mScarlet/PfRON3-mNeonGreen, parasite lines. White scale bars – 2 µm, yellow scale bars – 0.5 µm

To enable tracking of the MJ during invasion transgenic fluorescent *P. falciparum* lines were generated with mScarlet fused to either PfRON4 or PfAMA1. These parasite lines enabled the tracking of the MJ from the time it is established through to the completion of invasion, which involves sealing of the parasitophorous vacuole membrane (PVM) and restoration of the host cytoskeleton (**Supplementary Fig. 1a (iv)**). Additionally, dual-fluorescent lines were created by combining either the PfRON4-mScarlet or PfAMA1-mScarlet with PfRON3-mNeonGreen. These double-transgenic *P. falciparum* lines were designed to allow the release of the rhoptry neck (PfRON4) and bulb (PfRON3) components, or the micronemes (PfAMA1) and rhoptry bulb (PfRON3) components to be tracked either independently or simultaneously under different experimental conditions. Expression of these fusion proteins in transgenic parasites were validated with western blots using rat anti-HA antibodies against the HA tagged proteins (**Supplementary Fig. 2**).

The localisation of PfAMA1-mScarlet was validated using 3D Structured Illumination Microscopy (3D-SIM) in both late stage schizonts and free merozoites (**Fig. 1c**). In late stage schizonts PfRON3-mNeonGreen (Green) was shown to label the periphery of the rhoptry bulb membrane, consistent with that previously shown ^13^, while PfAMA1-mScarlet (magenta) was observed as a punctate morphology that does not collocate to the rhoptries, indicating its known localisation in the micronemes ^29^. Upon egress, PfAMA1 was released onto the merozoite membrane coating the exterior of the free parasite, in agreement with previous studies ^30^, and this was also observed for the PfAMA1-mScarlet lines which coat free merozoites. PfRON4-mScarlet was shown to localise to a single punctum at the tip of the PfRON3-mNeonGreen signal, indicating localisation to the rhoptry neck at the pre-invasion stage and prior to release (**Fig. 1d**).

### Dual fluorescent RON3-RON4 and RON3-AMA1 enable sequential tracking of invasion-related organelle release

Dual-labelled parasite lines (PfRON3-mNeonGreen/PfRON4-mScarlet and PfRON3-mNeonGreen/PfAMA1-mScarlet) were generated to enable the precise tracking of microneme, rhoptry neck and rhoptry bulb release. PfRON4 is a rhoptry neck localised protein, whereas PfRON3 localises to the rhoptry bulb, and when imaged together, it was possible to track the sequential release of these two rhoptry organelle compartments. Upon attachment of a merozoite to an uninfected erythrocyte, PfRON3-mNeongreen and PfRON4-mScarlet were seen to overlap during the initial deformations. Following the final deformation, during the recoil phase, PfRON4-mScarlet could be seen to insert into the host erythrocyte cytoskeleton at the localised region (see frame 00:17.70), which is subsequently cleared (as determined by SiR-actin fluorescence), whereas PfRON3-mNeonGreen remained extracellular. During internalisation PfRON4-mScarlet located to the aperture demarcating the MJ ^31,32^, whereas PfRON3-mNeonGreen remained punctate and translated to the intracellular space, indicating the apex of the invading parasite. The PfRON3-mNeonGreen signal diminished following the completion of internalisation signifying that the rhoptry bulb components are released during and after active invasion (**Fig 2a, Supplementary Movie 2**). By tracking the intensity of both PfRON3-mNeonGreen and PfRON4-mScarlet during the invasion process, it was also possible to plot the sequential release of these two compartments (**Fig 2c, Supplementary Movie 2**). The rhoptry organelles were segmented by creating a mask using the PfRON3-mNeonGreen signal and then measuring the mean intensity of both PfRON3-mNeonGreen and PfRON4-mScarlet within this region. Using this approach a clear temporal offset was measured between the release of the neck and bulb compartments of this club shaped organelle (**Fig. 2c**).

**Figure 2:**
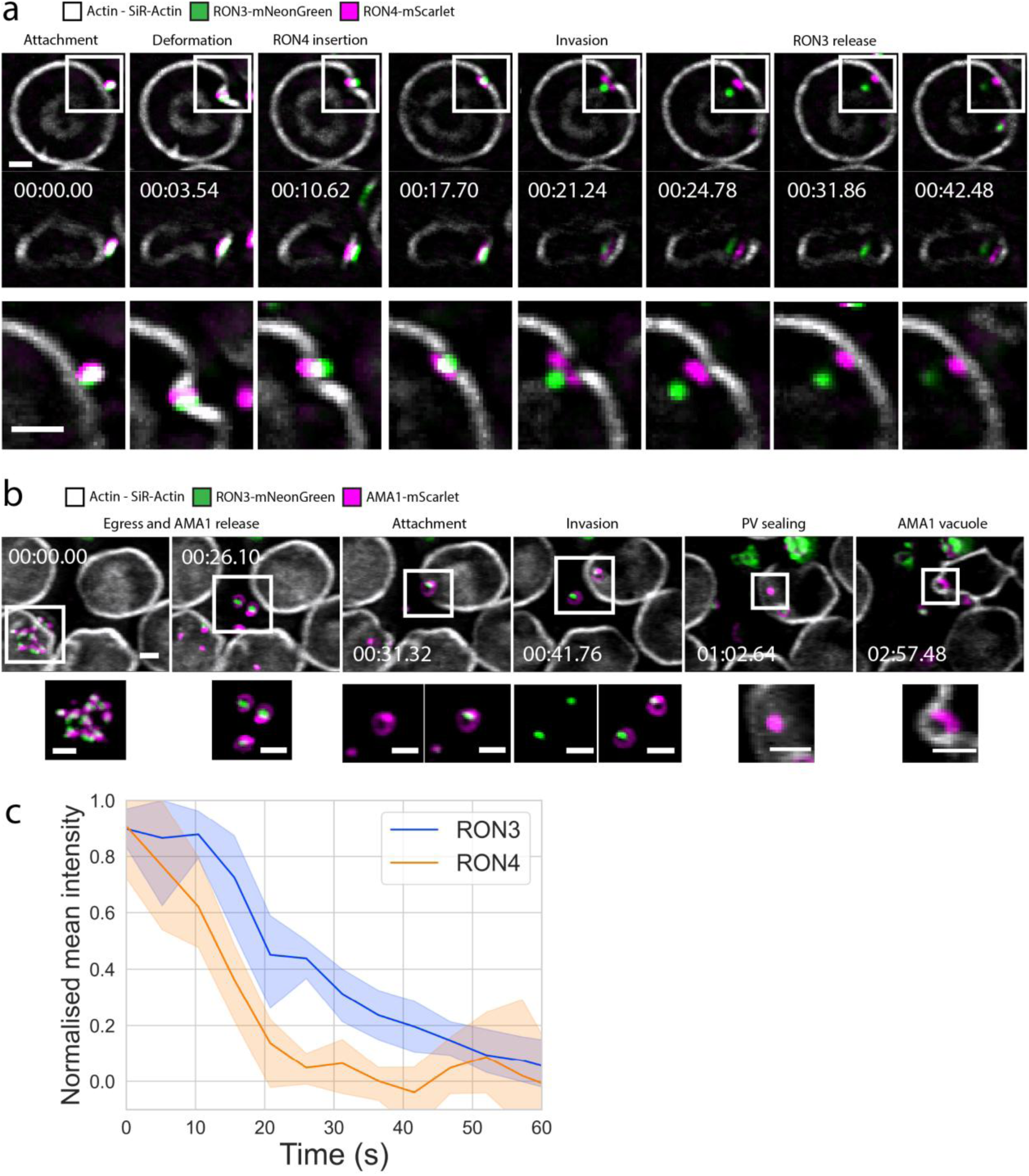
Dual fluorescent parasite lines to track subsequent release of invasion organelles. (a) Time-lapse sequence of dual PfRON4-mScarlet-PfRON3-mNeonGreen parasites during invasion imaged at a volume rate of one volume every 3.54 seconds. Scale bar – 2 µm. (b) Time-lapse sequence of invasion of dual PfAMA1-mScarlet-PfRON3-mNeonGreen fluorescent parasites at a volume rate of one volume every 3.54 seconds. Scale bar - 2µm. (c) Plot of the normalised mean intensity for both PfRON3-mNeonGreen and PfRON4-mScarlet during invasion. Intensities were measured within the segmented region of high PfRON3-mNeonGreen signal during invasion. Tracking the mean intensity of PfRON4-mScarlet withing the mNeonGreen region shows when RON4 is released relative to PfRON3 and then the subsequent release of PfRON3 is tracked through diminishing mNeonGreen intensity. Data plotted as the average of n=10 invasion events with 95% confidence intervals.

Combining PfAMA1-mScarlet with PfRON3-mNeonGreen, it was possible to track the release of micronemal proteins compared to those originating from the rhoptries. Just prior to egress the PfAMA1-mScarlet fluorescence was distinctly punctate as was the PfRON3-mNeonGreen signal. Immediately following egress PfAMA1-mScarlet was seen to more broadly label the exterior of the parasite indicating release to the parasite surface. When a merozoite attached to an uninfected erythrocyte, a bright PfAMA1-mScarlet region was seen at the apex that labelled the opening in the erythrocyte cytoskeleton during active invasion. The signal then localised to the erythrocyte surface at the point of sealing before localising to the PV, likely indicating the PfAMA1 on the parasite surface.

### The moving junction functions as a molecular sieve

It has previously been hypothesised that the MJ can act as a functional molecular sieve, which allows different lipids or proteins to be selectively incorporated into the newly formed parasitophorous vacuole membrane. Indeed, this has been shown to be the case particularly for *Toxoplasma gondii* ^22^. We have previously shown that blocking the formation of the MJ by inhibiting the interaction between PfAMA1-PfRON2 using R1 peptide ^33^, long lipid-based, membrane tethers or extensions can form from the site of interaction ^13^. Conversely, by inhibiting the parasite actomyosin motor through the disruption of actin polymerisation using cytochalasin D (Cyto D) ^34^, leads to intracellular protrusions ^13^. Using the dual fluorescent PfRON3-PfRON4 line, the incorporation of parasite protein to these protrusions or extensions relative to the established MJ, were tracked in real time (**Fig 3**). Upon Cyto D treatment, RON4-mScarlet was incorporated into the plane of the host cytoskeleton and this was then followed by ejection of PfRON3-mNeonGreen into a long protrusion within the intracellular space of the host cell (**Fig 3a**). This protrusion formed from the site of the MJ (shown by RON4) and grew over the course of minutes. When treated with R1 peptide, RON4-mScarlet could insert into the host cytoskeleton whilst PfRON3-mNeonGreen was ejected onto the surface of the erythrocyte (**Fig 3b**).

**Figure 3:**
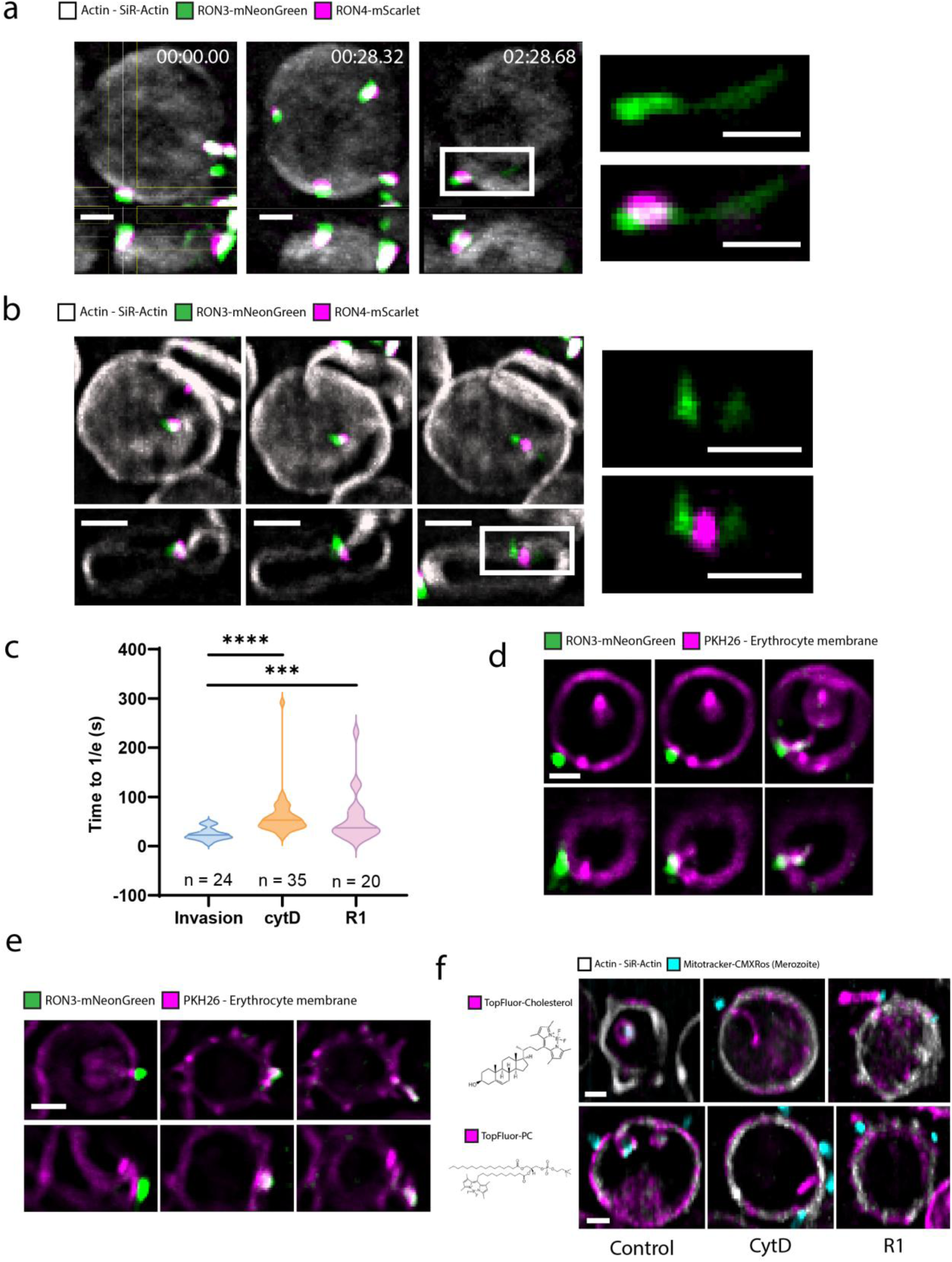
The moving junction functions as a molecular sieve during invasion. (a) Time-lapse snapshots of Cyto D inhibited PfRON4-mScarlet-PfRON3-mNeonGreen parasites during invasion. Images shown as a XY slice with YZ orthogonal view aligned with the interacting parasite. Inset shows PfRON3 and RON4 channels as PfRON3 is ejected into long intracellular protrusion. Scale bars – 2 µm. (b) Time-lapse snapshots of R1 peptide inhibited PfRON4-mScarlet-PfRON3-mNeonGreen parasites during attempted invasion event. Data presented as XY slice with YZ orthogonal section aligned to interacting parasite. Inset shows release of PfRON3 onto erythrocyte surface at the point of interaction denoted by insertion of RON4 signal. Scale bars – 2 µm. (c) Rate of PfRON3 release measured via 3D segmentation of PfRON3-mNeonGreen signal at interaction site and tracking of the total intensity of all voxels within segmented region. Following exponential fitting of mNeonGreen signal over time a constant, 1/e, was measured to denote the rate of rhoptry release. Significance determined by a two-tailed Mann-Whitney test. *** p < 0.001 (d) Representative extended section views, 1 µm section, of Cyto D inhibited parasite interacting with PKH26 labelled erythrocytes showing inward lipid labelled protrusions in combination with PfRON3-mNeonGreen single fluorescent parasites. Scale bar – 2 µm. (e) Representative XY and YZ extended section views, 1 µm section, of R1 peptide inhibited PfRON3-mNeonGreen parasites interacting with PKH26 labelled erythrocytes showing ejection of PfRON3 and formation of extracellular lipid extensions.

To compare the dynamics of rhoptry bulb release, we plotted the total intensity of PfRON3 fluorescence throughout the merozoite-erythrocyte interaction (Fig 3c). For Cyto D and R1 peptide conditions, interactions with a visible reduction in PfRON3-mNeonGreen fluorescence intensity were selected. Using one phase decay exponential curve fitting, the time between the maximum to 1/e (37%) of the intensity sum was extracted and used as a checkpoint for PfRON3 release. The PfRON3 release was found to be significantly slower in the presence of Cyto D (Median = 52.88 s, n = 35, N = 3) or R1 peptide (Median = 36.93 s, n = 20, N = 5) compared to invasion events occurring under normal conditions (Median = 22.64 s, n = 24, N = 4) (**Fig. 3c**).

To determine if the release of PfRON3-mNeonGreen was localised to membrane material, we labelled uninfected erythrocytes with the lipophilic dye PKH26. PKH26 provides a red fluorescent counterstain of lipid membranes and has previously been used to track PVM formation of invading merozoites ^13^. In the presence of Cyto D, PfRON3 could be seen to extend into the lipid protrusion at the site of attachment (**Fig 3d**). It was seen that the lipid protrusion into the cell formed before the release of PfRON3 into the erythrocyte suggesting that lipids extend beyond the MJ prior to the release of rhoptry bulb proteins, such as PfRON3. As only the host is labelled with PKH26, this indicates that host lipids can translate laterally through the MJ due to the PKH26 positive signal extend into the host cell. In contrast, when the MJ was not fully formed, lipid extensions have previously been shown when interacting merozoites are treated with R1 peptide ^13^. When imaging PfRON3-mNeonGreen parasites it was seen that the PfRON3 signal extended into the plane of the host membrane with overlapping PfRON3 and PKH26 signals (**Fig 3d**) and was also seen to be present in the extracellular lipid extensions following echinocytosis.

The capacity of the MJ to allow lipids, but not membrane bound proteins, to bypass the MJ was measured directly using conjugated fluorescent phospholipids and sterols. In the case of TF-cholesterol, signal was seen in the nascent PVM (control), lipid protrusions (Cyto D) and the lipid extensions (R1 peptide) indicating that cholesterol, and cholesterol rich regions, can translocate past the MJ (**Fig 3e**). Cholesterol is found to be enriched in ordered lipid domains, whereas phospholipids with neutral headgroups are more likely to be found in more disordered lipid domains. To determine if these more disordered domains are also able to bypass the MJ, invading parasites were imaged in the presence of erythrocytes labelled with TF-PC (Phosphatidylcholine) (**Fig 3e**) which would partition to more cholesterol deficient regions. In all cases, TF-PC labelling was like that of TF-cholesterol labelling the nascent PVM, and both lipid protrusions and extensions in treated conditions. This implies that the MJ was likely not selective for different lipids but likely delineates between lipid material and membrane associated protein content.

### The Ca^2+^ flux preceding parasite internalisation is not linked to breakdown of the host cytoskeleton

The function of the punctate Ca^2+^ flux during invasion remains enigmatic and various studies have hypothesised that it may be related to rhoptry discharge, moving junction formation and/ or breakdown of the host cytoskeleton ^3,9,35^. Previous studies employ a strategy whereby uninfected erythrocytes are labelled with the Ca^2+^ sensitive dye Fluo-4 AM, and it has been hypothesised that following the Rh5-basigin binding, a pore is formed at the merozoite-erythrocyte interface enabling Fluo-4 AM to flow into the parasite’s apical end where a punctate Ca^2+^ flux is observed^3,9,36^. To understand if this flux is occurring at the parasite apical end, we labelled the parasites with Fluo-4 AM and used erythrocytes labelled with SiR-Actin only. These parasites were able to interact and invade uninfected erythrocytes with all steps of invasion determined by LLSM (**Fig. 4a, Supplementary Movie 3**). After the final deformation and prior to erythrocyte membrane recoil, an increase in parasite Ca^2+^ was observed like that seen in the context of Fluo-4 AM labelled erythrocytes. This provides strong evidence the rise in Ca^2+^ is a functionally important second messenger involved in one of several steps known to occur at this stage of invasion (e.g. rhoptry discharge, motor engagement and cytoskeletal breakdown).

**Figure 4:**
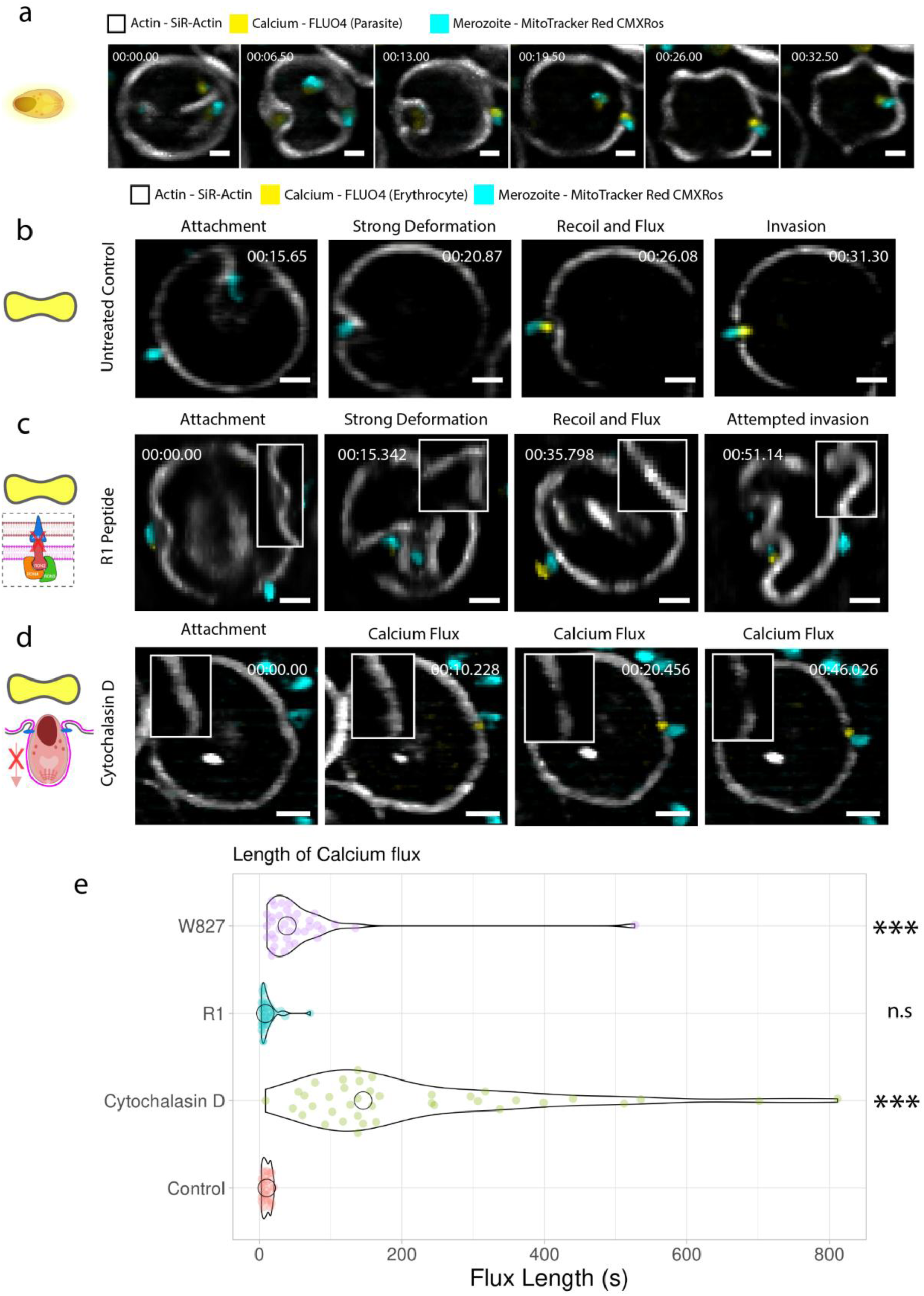
Ca2^+^ flux is functionally related to parasite motor engagement. (a) Representative images of LLSM data of invasion with FLUO-4 AM labelled merozoites. Data is represented as an extended slice view with a section thickness of 1 µm. Scale bar – 2 µm. (b-d) Representative images of Ca^2+^ flux and cytoskeletal breakdown during control invasion conditions, R1 peptide inhibition and Cyto D treatment, respectively, in the presence of FLUO-4 AM labelled erythrocytes. All images show as extended section view with a section thickness of 1 µm. All images were taken with LLSM with a volume rate of 1 volume per 3.54 s. Scale bars all 2 µm (e) Plot showing length of Ca^2+^ flux as measured from increase in parasite localised Ca^2+^ signal during invasion events. Length is measured from the first to last frames where increased Ca^2+^ signals were present. * p < 0.05, ** p < 0.01, *** p < 0.001; Randomisation test, n.s – no significance.

To determine the timing of the Ca^2+^ flux relative to rhoptry discharge and host actin breakdown, uninfected erythrocytes were labelled with both Fluo-4 AM and SiR-Actin and merozoites were able to interact and deform the host cell and reorientate. During the final deformation, a Ca^2+^ flux was observed, which preceded the breakdown of the host cytoskeleton (**Fig. 4b, Supplementary Movie 4**). Under conditions of R1 peptide inhibition, which blocks AMA1-RON2 binding, we were still able to measure the punctate Ca^2+^ flux and strong membrane deformations occurring upstream of MJ formation, however we did not observe any clearance of the host cytoskeleton (**Fig 4c, Supplementary Movie 4**). This is consistent with our previous data showing that, despite the strong membrane deformations, the same degree of negative Gaussian membrane curvature is not achieved implying an absence of cytoskeletal breakdown after R1 peptide treatment ^13^. By directly visualising host actin it was possible to observe that there is no breakdown of the host cytoskeleton in the absence of a functioning MJ despite a clear and consistent Ca^2+^ flux signal (**Fig 4c, Supplementary Movie 4**).

The effect of Cyto D, an actin polymerisation inhibitor, on both the Ca^2+^ flux and cytoskeletal breakdown during merozoite invasion was investigated next. Following egress, merozoites attached to other erythrocytes were able to induce a Ca^2+^ flux. Interestingly, on some occasions a clear breakdown in actin signal was observed at the point of apical alignment of the merozoite to host cytoskeleton (**Fig 4d, Supplementary Movie 4**). Cyto D inhibits not only the parasite’s ability to invade erythrocytes, but also its ability to induce pre-invasion deformations. Similar to control parasites, actin did not break down during the early phase of interaction but occurred after the punctate Ca^2+^ flux. This data points to an active disassembly process, likely involving the parasite’s MJ, with little evidence that the force of the parasite motor is contributing to mechanical disruption of the cytoskeleton during internalisation. Furthermore, during this interaction phase the Ca^2+^ flux was significantly prolonged compared to untreated or R1 peptide treated merozoites (**Fig 4e**).

To rule out an effect of Cyto D on the host cytoskeleton, we also tested *S*-W827^37,38^, a specific inhibitor of the parasite’s actomyosin motor that blocks the binding of profilin to actin ^37,38^. This inhibitor differs to Cyto D in that it specifically targets the parasite’s motor and does not affect erythrocyte actin polymerisation. As with Cyto D we were able to see active actin clearance at the point of attachment following Ca^2+^ flux. The temporal sequence of this matched that of untreated and Cyto D treated parasites in that actin broke down following Ca^2+^ flux indicating a well conserved sequence of steps. In addition, the length of the Ca^2+^ flux was prolonged significantly when compared with untreated or R1 treated parasites, suggesting Ca^2+^ homeostasis is impaired when the polymerisation of actin is inhibited. Taken together, these data indicate that the Ca^2+^ flux following PfRh5-basigin binding, does not mediate host cytoskeletal remodelling.

### Formation of the moving junction is coupled with cytoskeletal disruption

Following the surprising finding that actin clearance occurred in the absence of active invasion, we sought to determine the links between cytoskeletal breakdown and MJ formation. Up to now, experiments on the MJ have been largely limited to fixed imaging through immunofluorescence labelling or electron microscopy. AMA1 has been fluorescently tagged previously in both *P. falciparum* ^30^ and in *P. knowlesi* ^39^. To understand the temporal sequence of events of MJ formation, mScarlet labelled PfRON4 or PfAMA1 parasite lines were imaged with LLSM. Upon egress of PfAMA1-mScarlet merozoites, the fluorescent signal moved from a punctate state to marking the boundary of the parasite (**Supplementary Movie 5**). The membrane localised signal then aggregated at the invasion interface between parasite and erythrocyte and demarked the boundary between the cytoskeleton and parasite periphery throughout the invasion process. Following the completion of invasion, PfAMA1 remained localised to the parasite periphery, suggesting some PfAMA1 was retained on the parasite surface even after its internalisation and enclosure within the parasitophorous vacuole ^40^.

During the invasion of PfRON4-mScarlet merozoites, the signal remains punctate from egress through to the initialisation of invasion where RON4 was then seen to form a ring, which moves rearward as the parasite becomes internalised. RON4 fluorescence therefore demarks the boundary between the parasite and the host cytoskeleton throughout invasion, which is consistent with previous studies (**Fig 5a, Supplementary Movie 5**) ^31,41,42^. At the onset of internalisation, rhoptry-released RON4 translocates to the erythrocyte surface and integrates into the host cytoskeleton, evidenced by its planar insertion within the region of cleared actin. Within the temporal limits of our system, this occurred simultaneously with the Ca^2+^ flux. On some occasions, RON4 could be seen to translocate from the parasite to the erythrocyte cytoskeleton one frame earlier than the originating Ca^2+^ flux. This may suggest a role for Ca^2+^ flux downstream of rhoptry neck release. However, we were unable to adequately resolve the timing of these two events with the acquisition speed of 1 volume per 3.54 s. During early to mid-stage internalisation of the parasite, a clear ring of RON4 was visible, marking the boundary of actin clearance. Upon completion of invasion, RON4 remained punctate at the posterior end of the invading parasite where membrane sealing, and reformation of the host cytoskeleton takes place.

**Figure 5:**
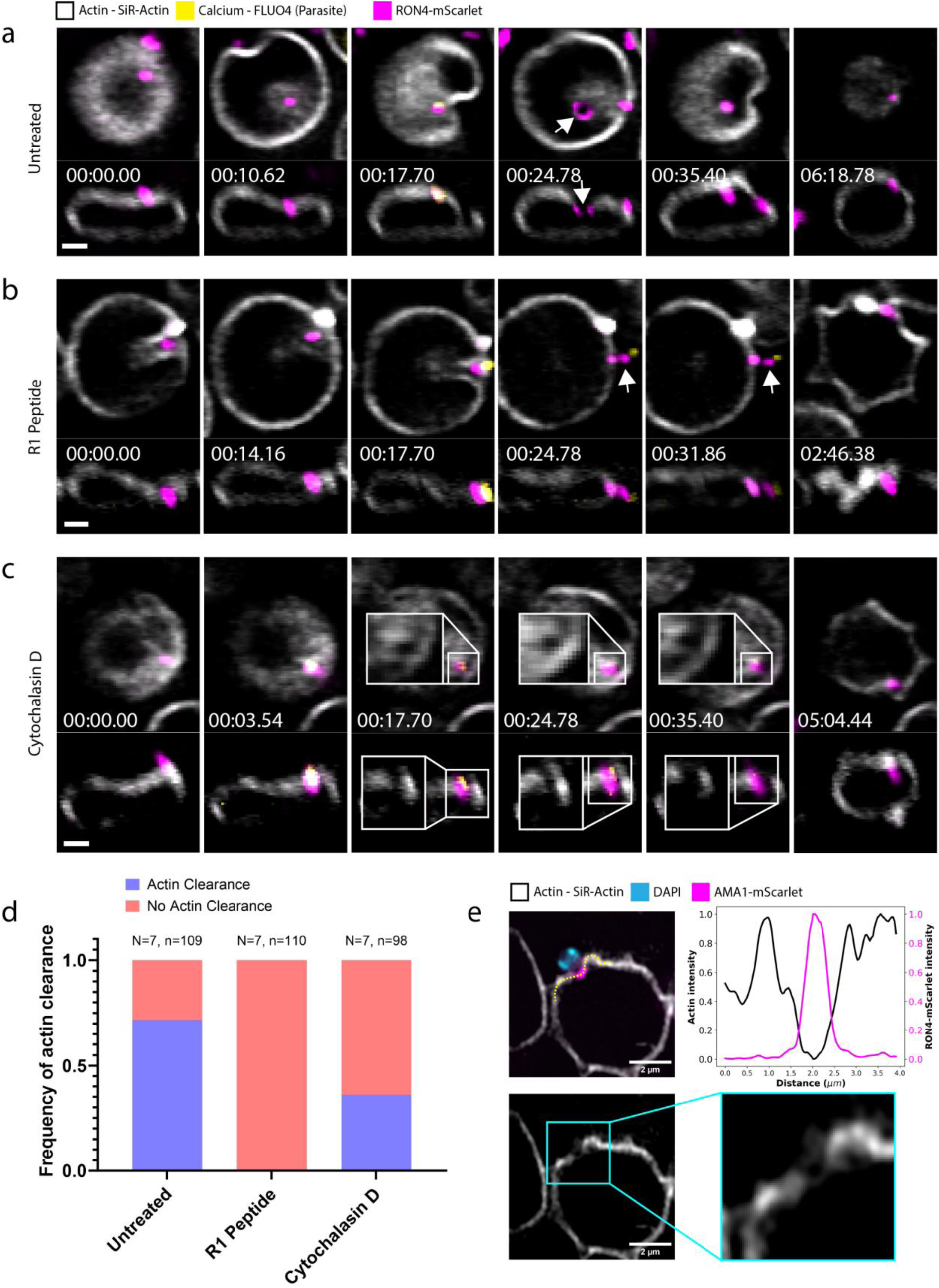
Fluorescent PfRON4 and PfAMA1 parasite lines demonstrate cytoskeletal breakdown is coupled to moving junction formation. (a) Representative images displaying key stages of invasion by PfRON4-mScarlet parasites including Ca^2+^ flux, MJ formation and cytoskeletal clearance. Images are displayed as slice view taken from volumetric LLSM data imaged at a volume rate of 1 volume per 3.54 seconds. (b) Time-lapse of R1 inhibited parasite interaction with Fluo-4 AM loaded erythrocyte displaying Ca^2+^ flux, PfRON4 insertion and subsequent PfRON4 splitting and echinocytosis. (c) Time-lapse of Cyto D inhibited parasite interactions with Fluo-4 AM loaded erythrocyte displaying Ca^2+^ flux, PfRON4 insertion, cytoskeletal breakdown and echinocytosis. (d) Frequency plot of host cytoskeletal breakdown determined for control invasion events, R1 peptide inhibition and Cyto D inhibition. Breakdown determined following positive Ca^2+^ flux signal and manually visualised in 3D or by a slice viewer. (e) Fixed immunofluorescence Airyscan confocal microscopy of Cyto D treated merozoite interacting with erythrocyte. Yellow dotted line indicates measured intensity profile in the plane of the host cytoskeleton and plotted as dual plot showing Actin and PfRON4-mScarlet intensities. Images show individual slice from 3D dataset. Scale bar – 2 µm.

Upon treatment with R1 peptide, where AMA1 binding of RON2 is blocked, PfRON4-mScarlet can be seen to insert into the erythrocyte cytoskeleton prior to the Ca^2+^ flux (**Fig 5b, Supplementary Movie 5**). Following the Ca^2+^ flux it could be seen that the PfRON4-mScarlet signal then split into two discrete puncta. One of these puncta was in the plane of the erythrocyte cytoskeleton, while the other was localised in the extracellular space, likely remaining in the parasite. These results suggest that the merozoites are still able to insert the RON complex to the host erythrocyte but are unable to incorporate the entirety of PfRON4 available for the invasion process.

When treated with Cyto D, the insertion of PfRON4-mScarlet signal into the plane of the erythrocyte cytoskeleton occurred where it remained in place up until the point of echinocytosis (**Fig 5c, Supplementary Movie 5**). In these experiments, actin was visibly cleared in 31% of cases where parasites had been treated with Cyto D, compared to 69% for invasions in untreated parasites. In contrast, there was no evidence of cytoskeletal breakdown when parasites had been treated with R1 peptide (**Fig 5d, Supplementary Movie 5**). We also investigated this using fixed immunofluorescence imaging of merozoites attempting to invade under Cyto D treatment conditions. This allowed us to interrogate the interaction region between erythrocyte and merozoite with higher spatial resolution using a Zeiss LSM 980 Airyscan confocal platform. Parasites were selected based on two criteria: 1) interacting with the erythrocyte perpendicular to the coverslip for maximum resolution in XY and 2) a visible embedding of the PfRON4-mScarlet signal into the host cytoskeleton (**Fig 5e and Supplementary Fig 3a**). With these criteria we measured the intensities of both PfRON4-mScarlet and actin along the axis of the erythrocyte cytoskeleton. A clear gap was measured in the host cell actin which was occupied by the embedded PfRON4-mScarlet signal. This was found to be consistent with merozoites also treated with *S*-W827 where a clear breakdown in the erythrocyte cytoskeleton was measurable with respect to PfRON4-mScarlet (**Supplementary Fig 3b**).

## DISCUSSION

Here, we quantitatively assessed the full sequence of erythrocyte invasion of *P. falciparum* merozoites using LLSM, together with a variety of fluorescent dyes and novel fluorescent parasite lines for labelling the erythrocyte membrane-cytoskeleton and parasite derived MJ complex, respectively. Through the application of these tools combined with inhibition at two distinct invasion checkpoints, we demonstrated that the cytoskeleton was cleared at the site of invasion downstream of the Ca^2+^ flux and that this was linked to the formation of the MJ as determined by the insertion of RON4 into the host cytoskeleton. This tool set offers a precise temporal understanding of the sequence of events during parasite mediated cytoskeletal breakdown, which has historically been limited to fixed points in time or to broad scale proteomic approaches.

It has not previously been possible to track the release of the different sub-compartments of the rhoptry organelles in real-time. This process has been often discretised into broad temporal events, such as initial merozoite contact, early, mid and late-stage invasion ^43^. This low-resolution view of the invasion sequence betrays the molecular complexity and functionally compartmentalised nature of the rhoptry organelles. By creating dual-fluorescent lines where the rhoptry organelle neck and bulb are labelled with different fluorescent proteins, we have demonstrated the temporal kinetics of this compartmentalised content release. Interestingly, the contents of the rhoptry bulb are not released until the parasite is actively invading the host cell. This release continues for some time following the formation of the PV suggesting an ongoing remodelling of the nascent PV immediately following the classical “completion” of invasion.

These new parasite lines have enabled us to also investigate the role the MJ plays in terms of molecular sorting and delineation between the host erythrocyte and newly formed PVM. It is known that for *T. gondii* that the MJ is capable of actively sorting molecular content that is incorporated to the nascent PVM ^22^. By utilizing the transmembrane rhoptry bulb component, PfRON3, it is possible to track the sequential release of the rhoptry organelle and the incorporation of parasite derived protein, into the newly formed PVM during invasion ^13^. When the MJ was formed, but invasion was inhibited through Cyto D treatment, PfRON3 was seen to extend into the large lipid protrusion extending into the host erythrocyte. This data shows that inhibited merozoites are still able to release the rhoptry bulb compartment, but transmembrane components remained restricted to the membrane material extending beyond the MJ. In the absence of a functioning MJ, through R1 peptide treatment, this release still occurred but was no longer restricted to the membrane material attached to the site of the PfRON4 signal. This indicates a role for the formation of the fully formed MJ to restrict the release of parasite derived material to the PVM. In contrast to this host lipid content was able to freely translate laterally via the MJ, whether intact or not, suggesting an active sorting mechanism of the MJ between lipid and protein material. This process was likely essential in maintaining the early replicative niche for the parasite in minimizing the localisation of parasite derived proteins to the exterior of the host cell (Figure 6b).

**Figure 6:**
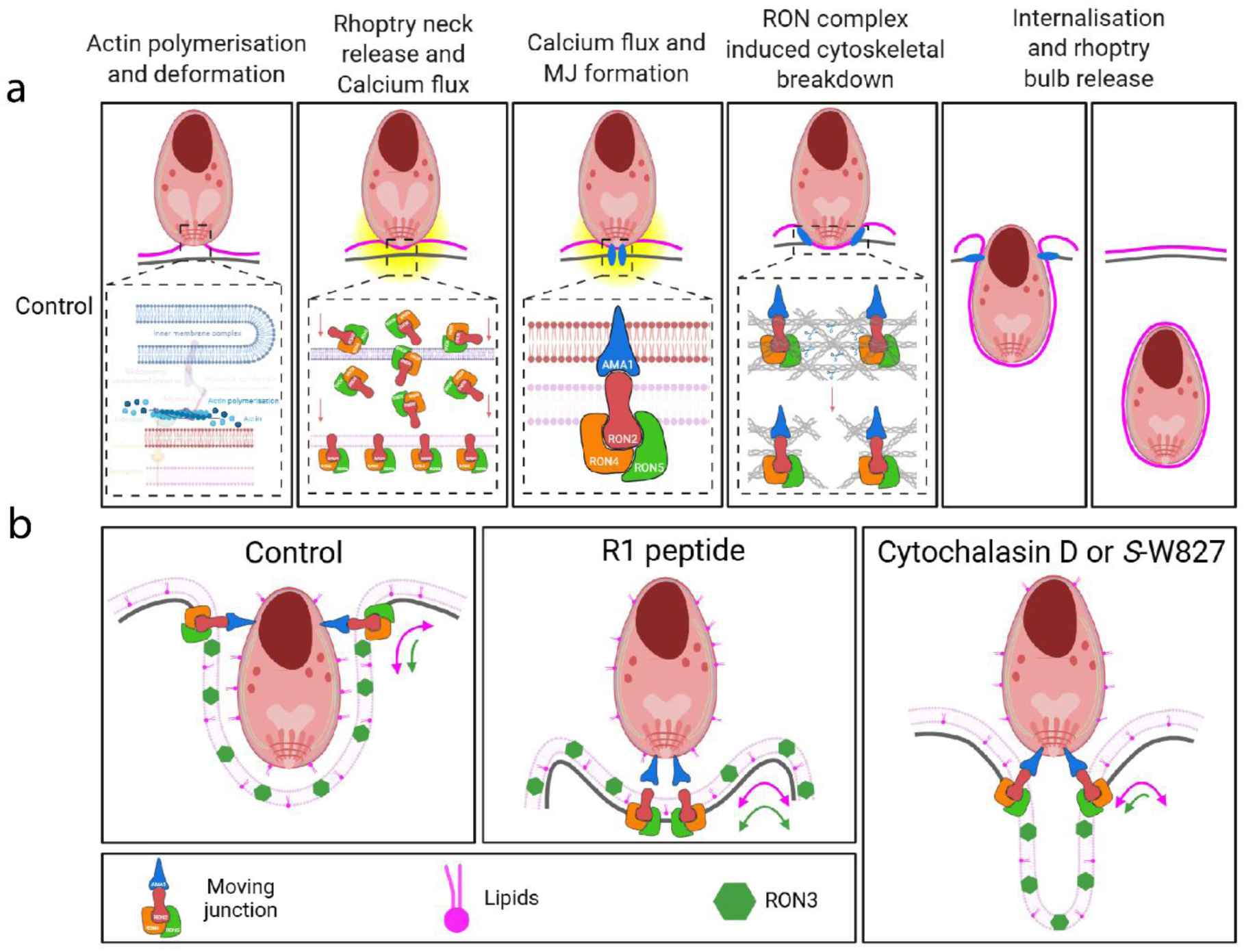
Schematic showing sequence of host cytoskeletal breakdown and its associated with formation. (a) Sequence of normal invasion conditions. Merozoites attached to the host erythrocyte surface and, following Ca^2+^ flux, insert the RON complex to bind AMA1. This step enables the active breakdown of the host cytoskeletal mesh network enabling successful invasion. (b) Schematic depicting a proposed molecular sieving mechanism when a functioning MJ is in place. Under normal invasion conditions proteins are restricted to the forming PVM, whereas lipids can freely translate beyond the MJ. Under R1 peptide inhibition, when the RON complex is inserted to the host cell but there is no cytoskeletal breakdown, proteins are also free to translate beyond the inserted RON complex due to no binding of AMA1-RON2. Under Cyto D, or *S*-W827, treatment parasites are able to attach, form a MJ and breakdown the host cytoskeletal mesh network, and ejected protein content is restricted to an inward lipidic protrusion in the host cell. This is also accompanied by a prolonged Ca^2+^ flux.

Previous studies show that *Plasmodium* invasion occurs normally in BAPTA-AM loaded erythrocytes ^3,44^. This led to the view that the PfRh5-basigin triggered Ca^2+^ flux does not directly contribute to local erythrocyte cytoskeletal breakdown nor echinocytosis during invasion ^3^. Our observations for a Ca^2+^ flux at the apical end of Fluo-4 AM labelled merozoites, which has identical characteristics to Fluo-4 AM loaded erythrocytes during invasion, strengthens this finding. In addition, we have shown that blocking the formation of the MJ using R1 peptide has no effect on the Ca^2+^ flux ^9,13^, but it blocks cytoskeletal breakdown. This confirms what we inferred from our previous findings that the cytoskeleton remains intact in the presence of R1 peptide because although parasites can generate substantial deformations, they do not achieve the high negative membrane curvature required for invasion ^13^. Taken together, the Ca^2+^ flux alone is unlikely to be responsible for mediating the breakdown of the host cytoskeleton, but the AMA1-RON2 interaction, required for MJ formation, is tightly linked to this process.

A prevailing hypothesis is that the Ca^2+^ flux arising from an uptake of extracellular Ca^2+^, is involved in rhoptry organelle secretion ^3^. This is based on the finding that removal of extracellular Ca^2+^ eliminates the Ca^2+^ flux and prevents both invasion and echinocytosis. However, in these conditions, parasites can still attach and deform the host cell ^3^. We have also uncovered an interesting finding that the Ca^2+^ concentration remains elevated when parasite actin polymerisation is inhibited. These results suggest that actin polymerisation, associated with the actomyosin motor, is crucial for the regulation of intracellular Ca^2+^ concentrations in the parasite apical region prior to internalisation.

The uptake of extracellular Ca^2+^ could produce rapid Ca^2+^ concentration rises reaching 100’s of micromolar beneath the parasite plasma membrane. Ca^2+^ homeostasis is essential for preventing sustained Ca^2+^ elevations from triggering a stress response. In addition, Ca^2+^ signals are typically organised into spatially and temporally restricted microdomains, which allow the selective activation of Ca^2+^ effectors ^45^. In *Plasmodium*, the regulation of Ca^2+^ signalling is achieved in part, through the uptake of Ca^2+^ by acidocalcisomes or by the endoplasmic reticulum (ER) through the action of a SERCA type Ca^2+^ ATPase (PfATP6) ^46^. Therefore, it is possible that the inhibition of actin polymerisation disrupts the spatial coupling between the ER and the plasma membrane and this in turn, disrupts the regulation of Ca^2+^. A loss of membrane and cytoskeletal tension could impair the restricted diffusion of Ca^2+^ and Ca^2+^ binding proteins. If so, the downstream activation of Ca^2+^ binding proteins, such as those associated with rhoptry secretion (e.g. Rasp2) require further analysis. Related to this, Ca^2+^ dependent protein kinases (CDPKs) have been identified in both *T. gondii* and *P. falciparum* ^47,48^. In *Plasmodium*, both PfCDPK1 and PfCDPK5 have been implicated in rhoptry neck secretion ^49^. PfCDPK1 is also implicated in the phosphorylation of proteins in the inner membrane complex and glideosome complex ^50^. The experimental system described here affords the opportunity to study RON4 release in combination with drugs inhibiting the uptake or release of ER Ca^2+^, or after the removal of extracellular Ca^2+^ ^51^, to address the precise order of events related to the Ca^2+^ flux and rhoptry neck secretion.

The appearance of an area of cytoskeletal clearance in the absence of a functioning actomyosin motor was surprising as, due to the diffraction limited nature of our data, this implies that even when the parasite was unable to deform the host erythrocyte, or engage its motor, it was still able to actively clear an area at the attachment site. The parasite’s motor was not only needed for active invasion, but also for pre-invasion deformations, which have been suggested to be involved in phosphorylation of the underlying cytoskeleton. To bypass the 80 nm hexagonal mesh of spectrin tetramers ^12^ the parasite has two options: either disassemble the associated junctional complexes at the invasion site or anchor its motor to these complexes and squeeze through relying on the lateral flexibility of the mesh ^52^. Our data point towards the former model and suggests that motor force is not responsible for mediating cytoskeletal clearance but may be needed to maintain it.

This data combined points to an integral role for the MJ in the critical step of host cytoskeletal breakdown and preservation of the PV during those important moments following a successful invasion (**Fig 6a**). While this data points towards a defining link between the MJ and the host-cytoskeleton, many questions remain. Exactly how does the MJ link into the host cytoskeleton? Structural studies are limited by the transient nature of this important structure and its inherent membrane bound localisation spanning not just the parasite plasma membrane but also that of the host cell. Studies in *T. gondii* have found various interacting molecular host partners which localise to the MJ during invasion such as CIN85, CD2AP, ALIX and TSG101 ^53^. CD2AP has also been identified as a modulator of host cytoskeleton for the human infecting parasite, *Cryptosporidium parvum (C. parvum)*^54^. In contrast to the host cells for *C. parvum* or *T. gondii*, CIN85 and CD2AP are not thought to be present in human erythrocytes so are unlikely to be involved in the formation of the MJ during *P. falciparum* invasion. In contrast, the ESCRT-I components ALIX and TSG101, are known to be present in human erythrocytes and are common markers of erythrocyte derived extracellular vesicles ^55–57^. In the case of *T. gondii*, a direct link was found between TgRON4 with ALIX, and TgRON5 with TSG101. The model presented suggests that ALIX and TSG101 could act as intermediates between the TgRON complex and underlying host cytoskeleton, in a ESCRT independent manner, due to the favourable binding sites present on the ALIX protein. While the examples in *T. gondii* have provided the first direct evidence of host-side partners for the RON complex, how these would then mechanistically lead to cytoskeletal disruptions remains unclear. It is possible that similar interactions could be present for PfRON4, PfRON5 and the erythrocyte cytoskeleton. These potential interactions could act as a platform for localised cytoskeletal disruption similar to that needed during cytokinetic abscission which is mediated by ESCRT proteins, such as ALIX and TSG101 ^58,59^. This could be achieved through indirect immunofluorescence of ALIX and TSG101 in invaded *P. falciparum* merozoites in the presence, and absence, of invasion inhibitors like Cyto D and R1 peptide. Ultimately, a deeper structural understanding of the RON complex, in particular the components RON4 and RON5, remains lacking due to challenges in recombinant expression of the full complex. *In-situ* methods could provide the necessary resolution, and molecular contrast, to determine the key interacting host protein partners. Imaging methods such as expansion microscopy ^60,61^ or *in situ* cryo-ET ^51,62^ may hold the keys to unlocking the secrets of this conserved, and essential, apicomplexan invasion machine.

## METHODS

### *P. falciparum* culture and synchronisation

Asexual stage 3D7 *P. falciparum* parasites were cultured in human O + erythrocytes at 4% hematocrit in RPMI1640 medium (Gibco) supplemented with 26 mM 4-(2-hydroxyethyl)piperazine-1-ethanesulfonic acid (HEPES), 50 mg/mL hypoxanthine, 20 mg/mL gentamicin, 0.2% NaHCO_3_, and 10% Albumax II (Gibco), as previously described ^63^. Use of human red blood cells from the Australian Red Cross (agreement 13-08VIC) was approved by the Walter and Eliza Hall Institute of Medical Research Human Research Ethics Committee (HREC86,17). The blood used for culturing *P. falciparum* parasites was obtained from human volunteers by the Australian Red Cross, with informed consent and was supplied deidentified. In brief, 30mL of culture was synchronised with two rounds of 5% sorbitol synchronisation. The culture medium was then discarded after centrifugation and cells were resuspended in culture medium in 5× volume of 5% sorbitol in a water bath at 37°C for 8 min. The sorbitol was then washed off and cells were resuspended in fresh culture medium. This synchronisation of the culture was further optimised by repeating this step after three days.

### Method for cloning of guides and parasite production

For HA-mScarlet tagging of *P. falciparum* RON4 gene (PlasmoDB accession PF3D7_1116000) the 5’ homology and codon optimised PfRON4 gene region was produced as a single contiguous gBlock from Integrated DNA Technologies (IDT) and inserted into the pR4HA-BSD vector using the restriction sites *Bgl* II/*Xma* I to create the pRON4HA-mScarlet-BSD plasmid. This vector has a pre-existing PfRON4 3’ homology region between the restriction sites *Eco* RI/*Kas* I. Guide oligos were designed to induce a double stranded break in PfRON4 at genomic position 3126 and InFusion cloned into pUF1-Cas9G using previously published methods ^64^.

For HA-mScarlet tagging of *P. falciparum* AMA1 gene (PlasmoDB accession PF3D7_1133400) an 884 bp 3’homology region was amplified from 3D7 gDNA with oligos DM778/779. The amplicon was cloned into pR4HA-mScarlet-BSD vector by the restriction sites *Eco*R I/*Kas* I to create pR4HA-mScarlet-BSD _AMA1 3’ vector. The 5’ homology and codon optimised AMA1 gene region was synthesised (Genscript) and cloned into pR4HA-mScarlet-BSD _AMA1 3’with *Not* I/*Xho* I restriction sites to produce pAMA1-HA-mScarlet-BSD vector. Guide oligos DM776/DM777 designed to induce a double stranded break in PfAMA1 at genomic position 2682 were InFusion cloned into pUF1-Cas9G using previously published methods ^64^.

RON3-HA-mNeonGreen parasites were transfected with either pAMA1-HA-mScarlet-BSD or pRON4HA-mScarlet-BSD along with the relevant pUF1-Cas9G guide vector as described previously ^6^ Transgenic parasites were recovered with 2.5 μg/ml blasticidin.

### Western blots

Synchronous 30 mL parasite cultures of PfRON3-mNeonGreen, PfRON4-mScarlet, AMA1-mScarlet, PfRON3-mNeonGreen-RON4-mScarlet, PfRON3-mNeonGreen-AMA1-mScarlet at 4% HCT and ≥5% schizont parasitaemia were pelleted (3,000 rpm/5 mins). The pellets were resuspended in 10X pellet volume of 0.15% saponin in 1X DPBS and incubated on ice for 10 mins before centrifugation (3,200 g/10 mins). The resulting pellets were wash washed 2-3X in ice-cold 0.15% saponin until supernatant was clear followed by a single wash with 1xDPBS (20,000 g/1 min). The parasite pellets were resuspended in 15X pellet volume of 4X reducing Laemmli SDS sample buffer (Thermo Scientific), sonicated for 10 cycles (30 secs on/off), reduced at 80 °C for 10 mins.

The samples were centrifugated (20,000 g/10 mins) before loading into a 4-12% NuPAGE™ Bis-Tris Mini Protein Gel (Invitrogen) and electrophoresed at 200 V along with Precision Plus Protein Dual Color Standards (Bio-Rad Laboratories). Electrophoresed proteins were transferred to nitrocellulose membrane using an iBlot™ Transfer Stack (Invitrogen) on the iBlot™ 2 Dry Blotting System (Invitrogen). The nitrocellulose membrane was blocked in 5% skim milk powder in 1X PBS with 0.1% Tween-20 and probed with horseradish peroxidase conjugated-rat anti-HA 3F10 antibody (1:500) (Roche), rabbit anti-aldolase R-759 antibody (1:2000) and horseradish peroxidase conjugated-goat anti-rabbit antibody (1:2,000) (Invitrogen). Blot was visualised with SuperSignal™ West Pico PLUS Chemiluminescent Substrate (Thermo Scientific) on the ChemiDoc Imaging System (Bio-Rad Laboratories). Aldolase was used as a loading control.

### Live-imaging with lattice light-sheet microscopy

To prepare the parasites for filming, the culture was loaded on LS columns attached to MACS MultiStand (Miltenyi Biotec) to isolate late-stage parasites. Imaging medium was prepared by adding 10 μM Trolox (Santa Cruz 53188-07-1) to the culture medium, with 0.25 mM CaCl_2_ and 5 mM sodium pyruvate (Gibco 11360070) also added when observing calcium flux. To compare effect of drug treatment on parasite invasion, either 1 μg/mL Cyto D (Sigma Aldrich C8273), 100 μg/mL R1 peptide (China Peptides), or 400 nM of S-W827 was added to the imaging medium. For host actin studies, erythrocytes were stained with 1 μM SiR-actin (Spirochrome SC001) in RPMI-HEPES. For membrane labelling, erythrocytes were stained with 1.5 μM Di-4-ANEPPDHQ (Invitrogen D36802) in RPMI-HEPES supplemented with 0.2% sodium bicarbonate. For calcium studies, erythrocytes were stained with 10 μM Fluo-4AM (Invitrogen F14201) in RPMI-HEPES supplemented with 5 mM sodium pyruvate. All staining mentioned above was done for 1 h at 37 °C. The stained erythrocytes were then washed three times and resuspended in imaging medium. For staining with PKH membrane dyes (Sigma Aldrich MINI26 and MINI67), erythrocytes were stained with 0.5 μM PKH dye in Diluent C at 1.5% hematocrit for 5 min at 37 °C. The stained erythrocytes were then washed once in medium with 10% serum, then washed twice in RPMI-HEPES before resuspending in imaging medium. For staining with TF lipid dyes (Avanti 810255 and 810281), erythrocytes were stained with 1 μM TF dye in malaria culture medium at 4% hematocrit overnight at 37 °C. The stained erythrocytes were then washed three times and resuspended in imaging medium. Purified schizonts were resuspended in culture medium and incubated with 10 nM Mitotracker Red CMXRos (Invitrogen M7512) for 30 min at 37 °C, 5% CO_2_. The stained schizonts were then pelleted and the supernatant removed before resuspending the schizonts in imaging medium. Before imaging, imaging medium was loaded to a well on an 8-well glass bottom plate (Ibidi 80807). Stained erythrocytes and stained schizonts were then added to each well and let settle.

All live-cell invasion imaging experiments were conducted using the Lattice Light-Sheet 7 (LLS7, Zeiss). Light from either a 488 nm, 560 nm, or 640 nm laser line, depending on the fluorophores used, illuminated the sample via a 13.3X 0.44 NA objective with a light sheet length of 30 µm and thickness of 1 µm. The resultant fluorescence was collected via a 44.83X, 1 NA detection objective and projected to a single camera via a 405/488/561/640 multi-band stop filter. A 250 x 250 x 30 µm was acquired with 726 slices providing an interval of 400 nm. Volume rates varied depending on the number of channels imaged, and the individual frame speed, between 2.8 – 7.7 seconds per volume. The overall resolution was measured to be 354 x 354 x 700 nm xyz as measured by the imaging of 100 nm multi-colour microspheres (Tetraspeck^TM^, Invitrogen) and were used to determine the optical setting of the aberration control which was set to 171.

### Lattice Light Sheet data processing

Raw data from the lattice light sheet microscope was deskewed and deconvolved prior to visualisation and analysis. As large regions of the image are either empty or contain no cells of interest, images were cropped on a region of interest (ROI) basis, to reduce processing time and save computational resources. ROIs were first defined on the maximum intensity projection of the data using Fiji ^65^, and then these regions passed to the napari-lattice pipeline ^66^. Data were deskewed with a skew angle of 30 degrees, followed by a coverglass transformation. A Richardson-Lucy deconvolution algorithm was applied to all data with 10-15 iterations and deconvolved using a point spread function defined by the imaging of 100 nm microspheres of different wavelengths.

### Structured Illumination Microscopy

Super-resolution three-dimensional structured illumination microscopy (3D SIM) was performed on the DeltaVision OMX-SR system (GE Healthcare) equipped with a ×60/1.42 NA PlanApo oil immersion objective (Olympus), sCMOS cameras, and 405, 488, 568, and 640 nm lasers, and 1.518 or 1.520 refractive index immersion oil. To image PfRON3, PfRON4 and AMA1 localisation on late-stage parasites, synchronised schizonts were labelled with DAPI. The 488 nm laser was used to excite mNeonGreen, 568 nm laser to excited either PfRON4-mScarlet or AMA1-mScarlet, and the 405 nm laser was used to excite DAPI. Structured Illumination image stacks were constructed from 15 raw images per plane (five phases, three angles) per color channel and a z-step size of 125 nm. Super-resolution reconstruction and color channel alignment were performed with softWoRx 7.0 (GE Healthcare).

### Percoll purification for fixed invasion imaging

30 mL parasite cultures (4% HCT) were resuspended and pelleted (1,900 RPM, 4 mins). Parasite pellet was resuspended in 4 mL supplemented RPMI (section P. falciparum culture) and dispensed over 6 mL Percoll density gradient (70% Percoll (Bio-Strategy, #GEHE17-0891-01), 7.8% 10X RPMI 1640 (Merck Life Science, #R4130-10X1L) 2.9% sorbitol (Merck Life Science, #S1876-5KG)) in a 15 mL conical tube (STEMCELL Technologies, #38009). The tube was centrifugated (3,500 RPM, 10 mins, 5 acceleration, 0 deacceleration). The dark layer containing late-stage parasites (trophozoites and schizonts) at the interface of the supplemented RPMI and Percoll density gradient was collected and dispensed into a 10 mL conical tube (Thermo Fisher Scientific, #LBSCT1203X) and washed with supplemented RPMI (1,900 RPM, 4 mins).

### Fixed merozoite invasion assay

The following assay was adapted as has been described ^31^. #1.5 13 mm round coverslips (Bio-Strategy, #CB00130RAC20MNZ0) were rinsed in 80% ethanol (in mili-Q water), placed in a 24-well tissue culture plate (Merck Life Science, #Z707805-72EA) and washed 3X with 0.22 µm-filtered 500 µL 1X PBS. The dried coverslips were coated with 500 µL 1% polyethylenimine (in mili-Q water; Merck Life Science; #P3143-100ML) at room temperature for ≥1 hour. The coverslips were washed 1X with 500 µL 1X PBS and left to dry in an opaque box for ≥30 mins before use. 30 mL PfRON4-mSc schizont cultures (4% HCT, ≥5% parasitaemia) treated with 30 nM ML10 (BEI Resources; MRT-0207065) overnight were Percoll purified. The resulting schizont pellet were resuspended in 20 mL RPMI 1640 with 10 µM E64 (Merck Life Science; #E3132) and incubated at normal culturing conditions for 1-2 hours until schizonts were almost all 48 hpi. Then, the schizonts were pelleted (1,900 RPM, 4 mins), resuspended in 2 mL supplemented RPMI (section P. falciparum culture), dispensed into a 5 mL syringe attached with a 1.2 µm filter, and the merozoites were isolated into a 10 mL conical tube. 500 µL of the merozoite suspension was dispensed into a 1.5 mL microcentrifuge tube containing 50 µL of 50% HCT red blood cell suspension in supplemented RPMI with compounds/peptides (final concentration: 0.1% DMSO or 1 µg/mL Cyto D or 100 µg/mL) and resuspended. The microcentrifuge tube was agitated at 37 °C (1,100RPM, 1 min 30 secs) in an Eppendorf ThermoMixer® C before fixation with 550 µL of 8% formaldehyde and 0.02% glutaraldehyde (final concentration 4% formaldehyde/0.01% glutaraldehyde). The cells were fixed on a roller for 30 mins at room temperature, then pelleted (3,000 RPM, 2 mins) and quenched/permeabilised by resuspension with 1 mL 0.1 M glycine + 0.1% Triton X-100 (in 1X PBS). The cells were pelleted and washed 1X with 1X PBS. The pellet was resuspended in 600 µL RPMI 1640 (Thermo Fisher Scientific, #72400047) with SiR-Actin dye (1:2000). 200 µL of the suspension was settled onto the polyethylenimine-coated coverslips for 20-30 mins at room temperature. The coverslips were removed from the 24-well tissue culture plate and egdes were dried on Kimwipes, then mounted on a microscope slide (ProSciTech, #AG00008032E01MNZ21) with 5 µL of VECTASHIELD® PLUS Antifade Mounting Medium with DAPI (Abacus dx, # VEH20002) and sealed with nail polish. Samples were mounted onto a Piezo stage and imaged on the Zeiss laser scanning microscope 980 with Airyscan 2 using the Plan-Apochromat 63X/1.40 Oil DIC objective with 405, 561 and 639 nm lasers. The following settings were used to acquire z-stacks: unidirectional frame scanning, 35 x 35 x 150 nm pixel size, 2 sampling rate, 0.84 µs pixel dwell time, 700 V (405 nm), 850 V (561 nm), 800 V (639 nm) detector gain, and 0.5% (405 nm), 5% (561 nm) and 2% (639 nm) laser power. Images were processed using [software].

### Quantitative analysis of RON3 and RON4 release

The analysis was performed on IMARIS 9.7.2 (Bitplane) with Tracking module, where PfRON3 signals were segmented and tracked surfaces were created with smoothing and background subtraction settings. The threshold was adjusted to similar values across all the data to include all PfRON3 signals that were visually distinguishable and 0.5 μm seed point value was used to split touching merozoites. Every PfRON3 surface that either gets internalised into a red blood cell or is released while in contact with a red blood cell was isolated and its track was monitored throughout the timelapse to ensure accurate tracking. The track was then cut such that the first frame of the track is the beginning of the merozoite contact on the red blood cell. The ‘Intensity Sum’ value of PfRON3 fluorescence was extracted from the isolated surface track and exported to Microsoft Excel. The values were then plotted up to 300 s and a one-phase decay exponential curve was fitted using GraphPad Prism 9.5.0. Any event with low goodness of fit (R squared less than 0.7) on the curve fitting was disqualified, then the Tau value (time elapsed when the curve reaches 1/e or 37% of the maximum value) from each qualified event was collected and used as a measure for the time taken between initial contact and PfRON3 release. For statistical analysis, two-tailed Mann-Whitney test was performed using GraphPad Prism 9.5.0.

### RON4 and Actin intensity line plot analysis

Line intensity profiles were manually drawn for invading merozoites under Cyto D treatment using the freehand drawing tool in Fiji/ImageJ. Intensity data was then plotted for both PfRON4-mScarlet and SiR-Actin along the contour of the erythrocyte cytoskeleton. These data were normalised to the maximum intensity for each channel to show the relative change in intensity along the line profile.

### Reporting summary

Further information on research design is available in the Nature Research Reporting Summary linked to this article.

## Data availability

A subset of data used in this study is available on Bio archive or zenodo. Source data are provided with this paper.

## Code availability

All customised ImageJ macros or other forms of data code used during this study are available upon reasonable request.

## ACKNOWLEDGEMENTS

We thank Kirsten Elgass from Zeiss Germany for helping to support the establishment of the Lattice Lightsheet 7 at WEHI. We acknowledge the Australian Red Cross Blood Service for providing blood. The authors gratefully acknowledge the WEHI Centre for Dynamic Imaging for their support and assistance in this work. This work was supported by an EMBO Long Term Fellowship ALTF 793-2016 and Sir Henry Wellcome Fellowship 206515_Z_17_Z to M.P. This work was also supported by grants from the National Health and Medical Research Council (NHMRC), APP1177431 to D.M, and NHMRC APPs 1121178, 1092789 and 1117288 to A.F.C, and 2012271 to K.L.R.

## CONTRIBUTIONS

N.D.G., C.E. and D.B.L. performed the LLSM experiments. C.E., N.D.G. and D.B.L. optimised sample preparation protocols for live cell imaging. C.E., and D.B.L prepared parasites. D.B.L. performed the immunofluorescence assay and prepared the western blots. A.D. and D.M. generated the mScarlet-tagged RON4 and AMA1 parasite lines. M.M performed 3D SIM. N.D.G., P.R and L.W.W. established the data processing and analysis pipelines. N.D.G., C.E., D.B.L and K.L.R., performed data analysis. W.N and B. E. S. synthesised compounds used in this study. N.D.G., K.L.R., C.E., and A.F.C. conceived and designed experiments. N.D.G., K.L.R., C.J.T., and A.F.C., supervised the work. N.D.G., C.E., and K.L.R. wrote the paper.

