## Supplementary figures 1-4 for "Formation of the moving junction is the nexus for host cytoskeletal remodelling during *Plasmodium falciparum* invasion of human erythrocytes"

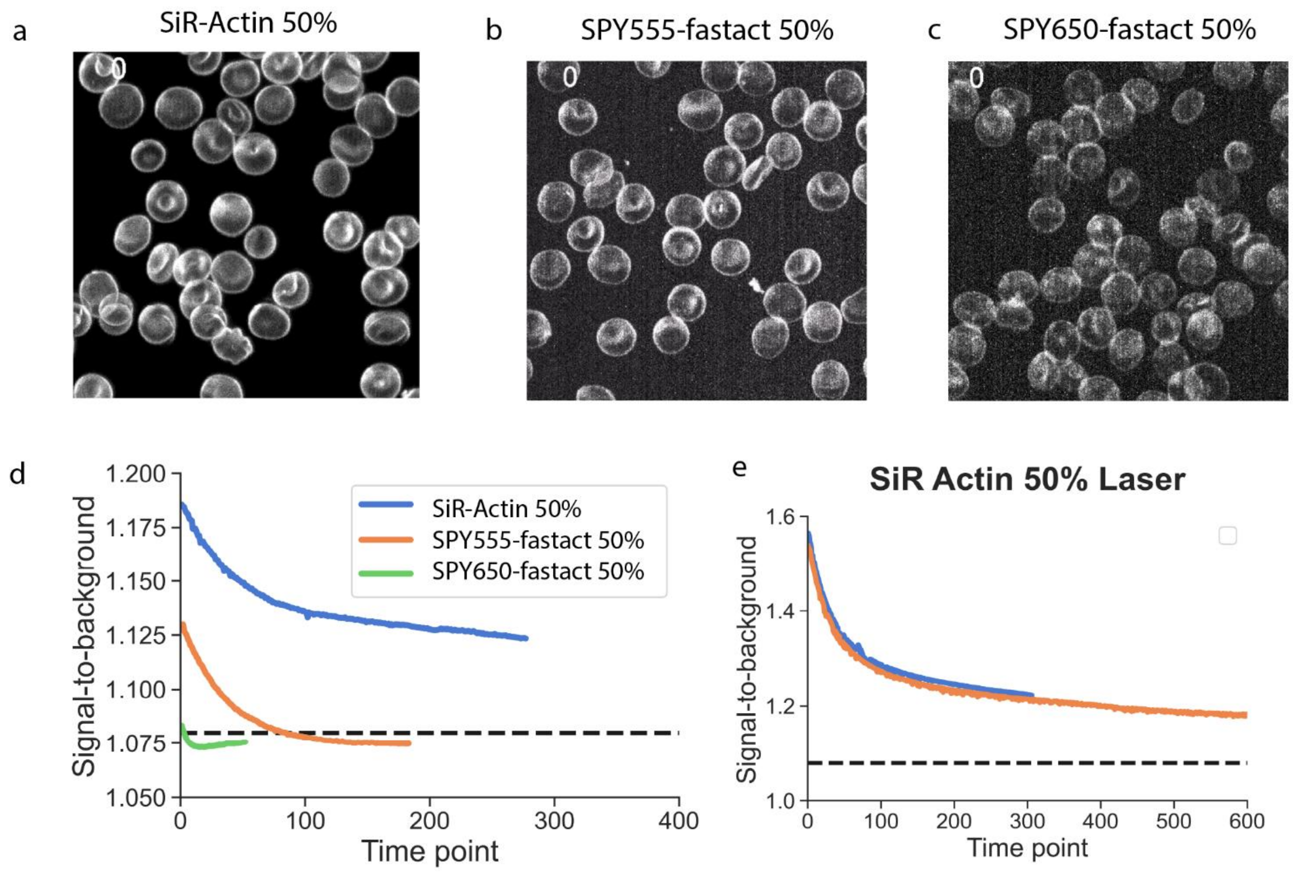


**Supplementary Figure 1: (a)** Erythrocytes labelled with SiR-Actin imaged by Lattice Light-Sheet Microscopy (LLSM) at 50% laser power with a 2 ms exposure time, **(b)** Erythrocytes labelled with SPY555-Fastact imaged by Lattice Light-Sheet Microscopy (LLSM) at 50% laser power with a 2 ms exposure time, **(c)** Erythrocytes labelled with SPY650-Fastact imaged by Lattice Light-Sheet Microscopy (LLSM) at 50% laser power with a 2 ms exposure time, **(d)** Signal-to-background ratios for a field of erythrocytes labelled with different actin markers plotted over consecutive imaging frames. 50% laser power and a 2 ms exposure time was used for all dyes. The dotted line shows the minimally detectable limited for sufficient measurements of erythrocyte actin over time. The SiR-Actin probe was found to provide sufficient signal to background over 200-300 imaging frames as necessitated by a standard LLSM invasion imaging experiment, (**e**) Signal-to-background ratios for a field of erythrocytes labelled with SiR-Actin imaged over 300 and 600 consecutive volumes with a frame exposure time of 2 ms. A steady state signal above the minimal detectable signal-to-background was found following a high peak signal.

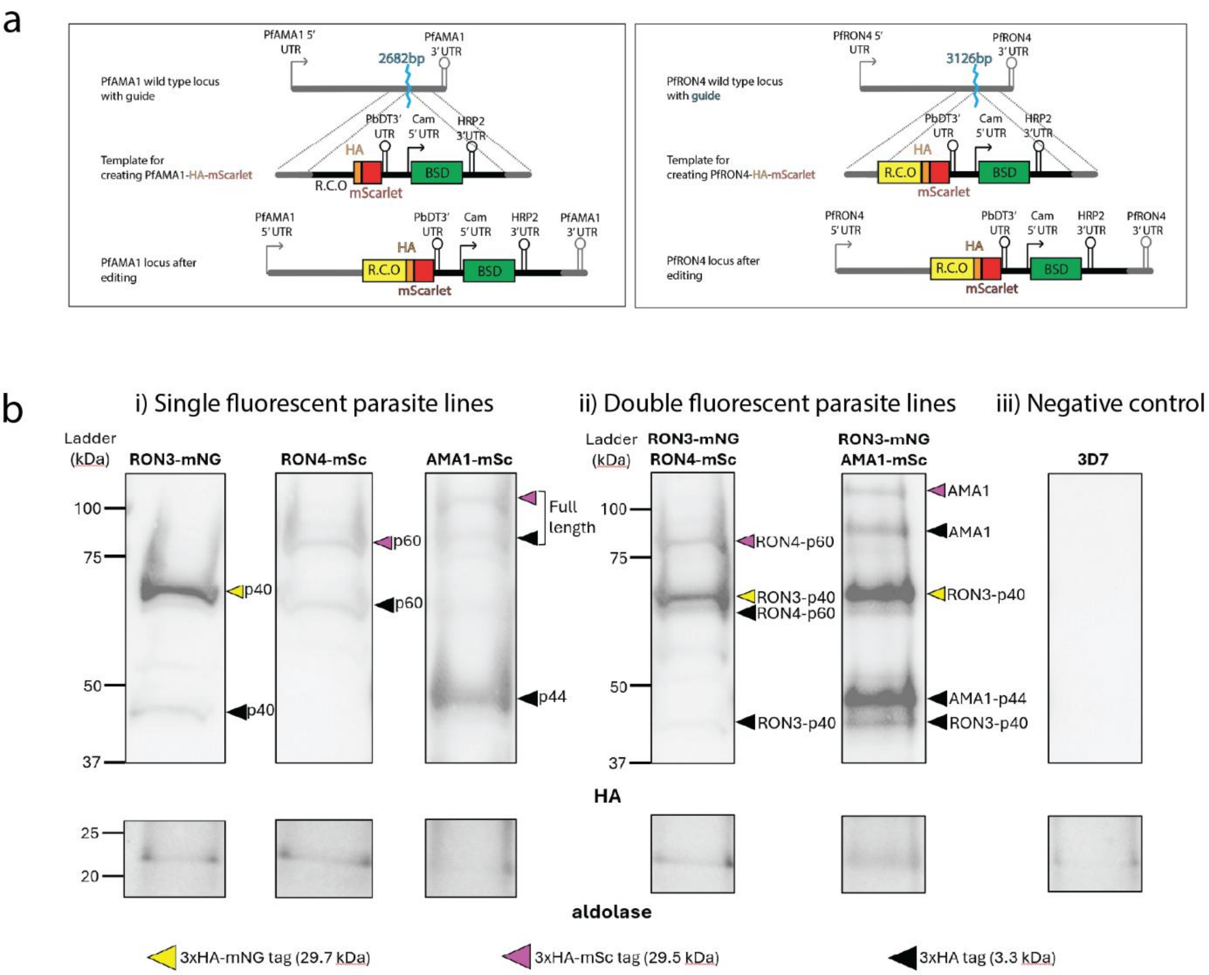


**Supplementary Figure 2: (a)** CRISPR-Cas9 targeting strategy to insert HA-mScarlet epitope tags at the 3' end of coding regions of PfAMA1 and PfRON4. BSD drug resistance marker allows for selection of successfully modified transgenic parasites. **(b(i))** Western blot of 3D7 parasites transfected with PfRON3-HA-mNeonGreen (mNG), PfRON4-HA-mScarlet (mSc) and PfAMA1-HA-mSc with proteins harvested from schizonts. In RON3-mNG, the anti HA antibody detected the cleaved C-terminal RON3 fragment (40 kDa) at ~43 kDa (HA tag) or ~70 kDa (HA-mNG tag); In RON4-mSc, the cleaved C-terminal RON4 fragment (60 kDa) at ~63 kDa (HA tag) or ~90 kDa (HA-mSc tag); In AMA1-mSc, the full length AMA1 (83 kDa) at ~87 kDa (HA tag) or ~113 kDa (HA-mSc tag), cleaved N-terminal AMA1 fragment (44 kDa) at ~47 kDa (HA tag). **(b(ii))** Western blot of transgenic RON3-mNG parasites transfected with PfRON4-HA-mSc and PfAMA1-HA-mSc with proteins harvested from schizonts. The HA antibody detected the same HA- or HA-mNG or HA-mSc-tagged bands as previously in both RON3-mNG-RON4-mSc and RON3-mNG-AMA1-mSc. **(b(iii))** No bands were detected in the negative control 3D7 parasite line. Aldolase was used as a loading control (~24 kDa).

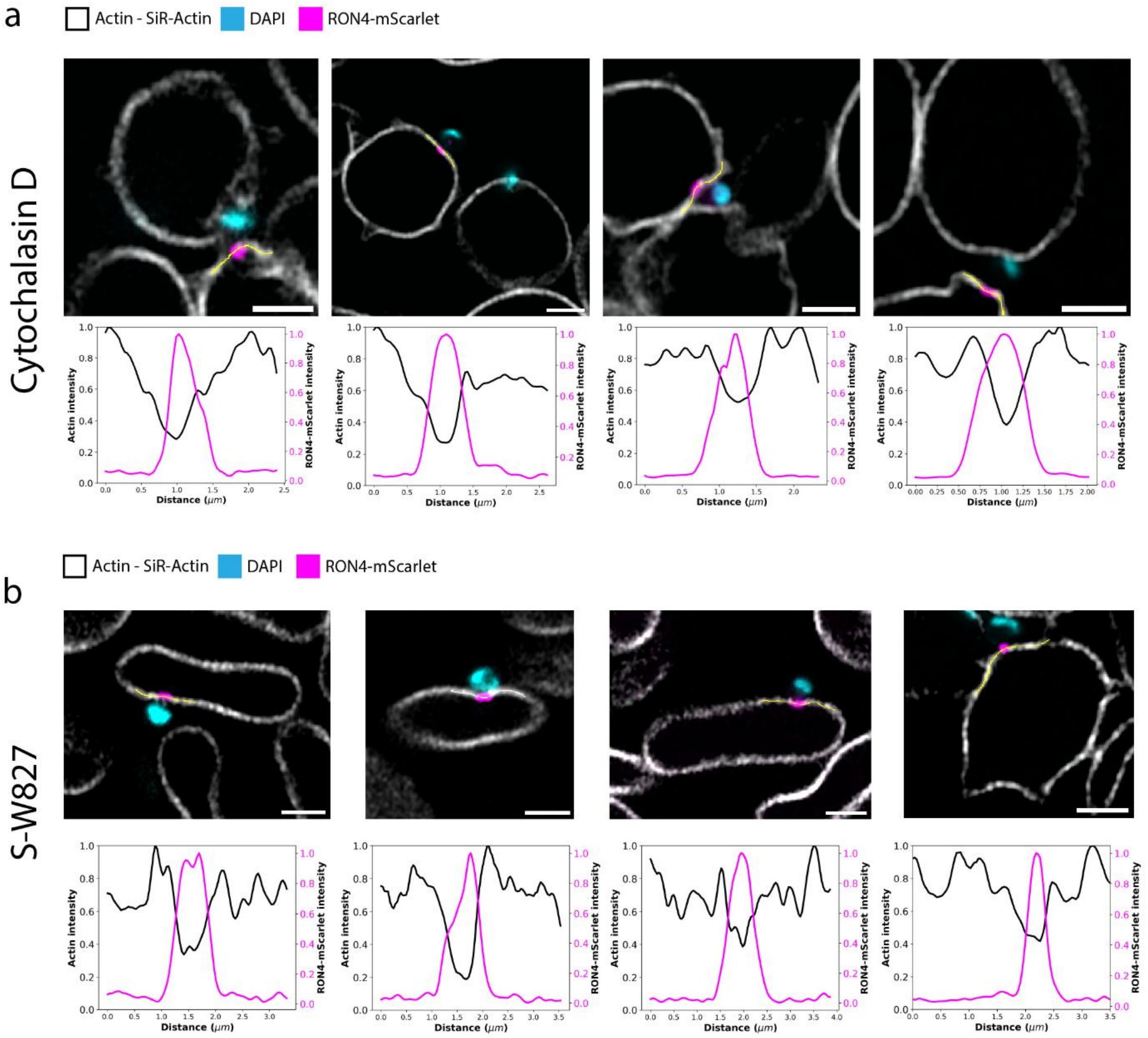


**Supplementary Figure 3: (a)** Examples of cytoskeletal breakdown following merozoite attachment to uninfected erythrocytes upon cytochalasin D treatment. Graphs represent the normalised intensities of Actin (Black lines) and PfRON4-mScarlet (Magenta lines) related to the plotted regions (yellow lines) on the associated images. Images are of fixed immunofluorescence prepared samples imaged using a Zeiss LSM 980 with the Airyscan mode, **(b)** Examples of cytoskeletal breakdown following merozoite attachment to uninfected erythrocytes upon *S*-W827 treatment. Graphs represent the normalised intensities of Actin (Black lines) and PfRON4-mScarlet (Magenta lines) related to the plotted regions (yellow lines) on the associated images. Images are of fixed immunofluorescence prepared samples imaged using a Zeiss LSM 980 with the Airyscan mode

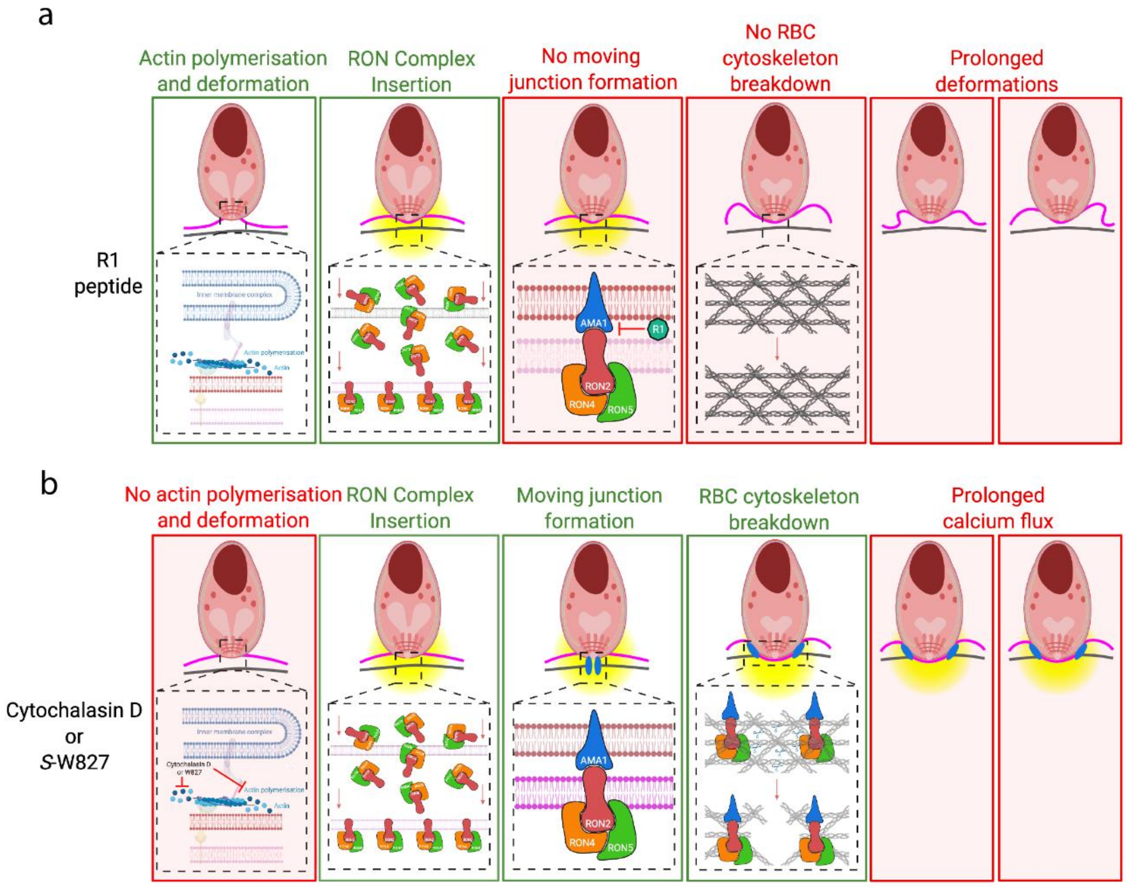


**Supplementary Figure 4: (a)** Sequence of events upon inhibition of invasion following R1 peptide treatment. Following attachment to an uninfected erythrocyte the RON complex is inserted to the host cell but without any associated cytoskeletal breakdown. **(b)** Sequence of invasion steps of parasites inhibited with compounds targeting the parasite's acto-myosin motor. Parasites are able to attach, form a TJ and breakdown the host cytoskeletal mesh network. This is accompanied by a prolonged calcium flux.
